# Genomic characterization of SARS-CoV-2 from Islamabad, Pakistan by Rapid Nanopore sequencing

**DOI:** 10.1101/2022.02.17.480826

**Authors:** Nazish Badar, Aamer Ikram, Muhammad Salman, Massab Umair, Zaira Rehman, Abdul Ahad, Hamza Ahmed Mirza, Masroor Alam, Nayab mehmood, Uzma Bashir

**Affiliations:** National Institute of Health, Islamabad, Pakistan; World Health Organization

**Keywords:** SARS-CoV-2, Pakistan, COVID-19

## Abstract

Since the start of COVID-19 pandemic, Pakistan has experienced four waves of pandemic. The fourth wave ended in October, 2021 while the fifth wave of pandemic starts in January, 2022. The data regarding the circulating strains after the fourth wave of pandemic from Pakistan is not available. The current study explore the genomic diversity of SARS-CoV-2 after fourth wave and before fifth wave of pandemic through whole genome sequencing. The results showed the circulation of different strains of SARS-CoV-2 during November-December, 2021. We have Omicron BA.1 (n=4), Lineage A (n=2) and delta AY.27 (n=1) variants of SARS-CoV-2 in the population of Islamabad. All the isolates harbors characteristics mutations of omicron and delta variant in the genome. The lineage A isolate harbors a nine amino acid (68-76) and a ten amino acid (679-688) deletion in the genome. The circulation of omicron in the population before the fifth wave of pandemic and subsequent upsurges of COVID-19 positive cases in Pakistan highlights the importance of genomic surveillance.

## Introduction

The emergence of SARS-CoV-2 in late 2019 and its variants have been responsible for more than 5.6 million deaths globally [1]. Over the period of two years the virus has incorporated many changes in its genomes. The first dominating mutation identified in the original Wuhan-Hu-1 virus was D614G that makes the virus more transmissible by increasing the binding affinity of spike protein with ACE2 receptor [2, 3]. During the late half of 2020 the virus has incorporated many new mutations in the genome leading to the emergence of various variants of concern (alpha, beta, gamma, and delta) and variants of interest. The emergence of SARS-CoV-2 variants has maintained the global transmissibility of SARS-CoV-2 even with rigorous vaccination drive in various countries [4]. Recently, in November 2021, a variant named Omicron emerged in South Africa and rapidly spread in more than 100 countries [5]. The omicron variant has caused successive epidemics of infection and reinfection of SARS-CoV-2 in countries with significantly vaccination coverage.

Pakistan has also been affected by COVID-19 pandemic and witnessed upsurges/waves of COVID-19 cases. The first cases of COVID-19 was reported in February, 2020. After wards four epidemic waves have struck in February, 2020, October, 2020, March, 2021 and July, 2021 respectively. Till November, 2021 the COVID-19 positivity rate in Pakistan has declined to 0.5% [6]. The continued genomic surveillance of circulating lineages of SARS-CoV-2 within population is crucial in order to forsee the emergence of new variants. The genomic surveillance has revealed the dominating lineages of SARS-CoV-2 during each epidemic wave but the data regarding the circulating lineages after the fourth wave of pandemic is not available. Hence, the current study has been conducted to understand what strains are circulating in Pakistan that were responsible for flattening the pandemic curve in Pakistan.

## Materials and methods

### Sample collection

During the months of November and December 2021, oropharyngeal swab samples were collected from 3500 suspected COVID-19 patients at National Institute of Health. The National Institute of Health’s Internal Review Board accepted the study design, and the datasets were anonymised and free of personally identifiable information.

### RNA Extraction and RT-qPCR

Viral RNA was isolated from the samples using MagMAX™ Viral/Pathogen Nucleic Acid Isolation Kit and KingFisher ™ Flex Purification System (ThermoFisher Scientific, US). The TaqPathTM COVID-19 CE-IVD RT-PCR kit (ThermoFisher Scientific, Waltham, US) was used to identify the presence of SARS-CoV-2, which targets three genes (ORF1ab, N, and S).

### Library preparation for MinION sequencing technology

The extracted RNA from SARS-CoV-2 positive samples (ct value < 30) were reverse transcribed with Luna script cDNA master mix and utilized as primary input for overlapping tiling PCR reactions spanning the viral genome with New England Biolabs Q5 High-Fidelity 2 Master Mix (M0492L) (primers provided in Supplementary Table S1). The ARTIC Network amplicon sequencing procedure v2 and the v3 primer pools were used to create amplicon pools (Quick, 2020). PCR product pools were quantified using a Promega Quantus fluorometer after purification.

The ligation sequencing kit from ONT was used to prepare the libraries (SQK-LSK109). The native barcoding expansion 96 kit was used for multiplexing (EXP-NBD114). DNA repair and end-prep were conducted with NEBNext Ultra II End-Repair/dA-tailing (New England Biolabs) with 1000 fmol of input cDNA and incubation periods were raised to 30 minutes at 20°C, followed by 30 minutes at 65°C, according to the manufacturer’s recommendations. For the barcoding reaction, 200 fmol of input cDNA were incubated for 20 minutes at room temperature (15–25°C) with the native barcodes and Blunt/TA Ligase Master Mix (New England Biolabs), followed by 10 minutes at 65°C.

For equal representation in the final library combination, equal numbers of indexed products were blended. To increase the efficiency of barcode ligation by using smaller pieces. On each minION flowcell, up to ten clinical samples (90 fmol/sample) were multiplexed, with a no-template control (NTC) processed in each pooled library phase. A DNA library was placed onto a MinION Spot On flowcell that had been primed (R9.4). Samples were sequenced for 72 hours using FLO-MIN106 flowcells on MinION MK1b sequencing equipment with high sequence basecalling enabled using MinKNOW (Version 4.2.8, Oxford Nanopore Technologies).

The BaseStack software platform [7] was used to create consensus genomes using a modified ARTIC network pipeline v1.0.0. Read length filtering and reference alignment to the Wuhan-Hu-1 genome were used to test variant polishing in Nanopolish v0.13.2, Medaka v0.11.5, and samtools 1.9. (GenBank accession number MN908947.3). The lineage assignment was done through PANGOLIN v2.1.7 [8]. The mutation profile analysis was performed through Nextstrain [9].

### Genomic dataset and Phylogenetic classification

For phylogenetic tree NCBI BLAST search of the study isolates were performed in order to get closely related sequence of SARS-CoV-2. Additionally sequences from neighboring countries were also included in the analysis. For multiple sequence alignment MAFFT software was used. For substitution model prediction jModelTest was used, Maximum Likelihood (ML) phylogenetic tree was build using IQtree (http://www.iqtree.org/). Tree was rooted with reference, Wuhan SARS-CoV-2 (hCoV-19/Wuhan/IME-WH01/2019). The tree was edited and visualized using Figtree software (http://tree.bio.ed.ac.uk/software/figtree/).

## Results

Between November 1 and December 31, 2021, a total of 1500 samples were tested positive for the presence of SARS-CoV-2 at the National Institute of Health’s virology department using the TaqPathTM Real-time RT-PCR kit (ThermoFisher Scientific, Waltham, US). Representative samples (n= 10) with Ct values <30 were selected for Nano pore sequencing during the study period. Due to a failed QC following the cDNA enrichment phase, 03 samples were not processed further. All the seven study participants were belonged to Islamabad. The selected subjects ranged in age group from 3-35 years old with median age of 27 years. The male to female ratio was 3:4.

### Phylogenetic profiling

Phylogenetic analysis of 7 full length SARS-CoV-2 genomes from this study and 101 global isolates was conducted (Fig 1). Of the 4 genomes sequenced of this study, make one clustered with Omicron sequences from USA and closely related with Scotland, Netherlands, and Ireland. Two genomic sequence of this study make separate clustered with Lineage A sequences reported from Pakistan and having nine amino acid deletion and closely related with Shanghai sequence (Fig 1). One genomes sequence of this study, make one clustered with delta sequences from USA. These seven sequence strains were all from Islamabad (S1 Table), suggesting that the viral strains circulating in the city were predominantly Lineage A, Delta and Omicron. The mean pairwise genetic distance between our sequences; sequences previously deposited from Pakistan, and sequences from USA, China, and Netherlands was found to be 0.00, indicating phylogenetic relatedness between the genomes.

**Figure 1:**
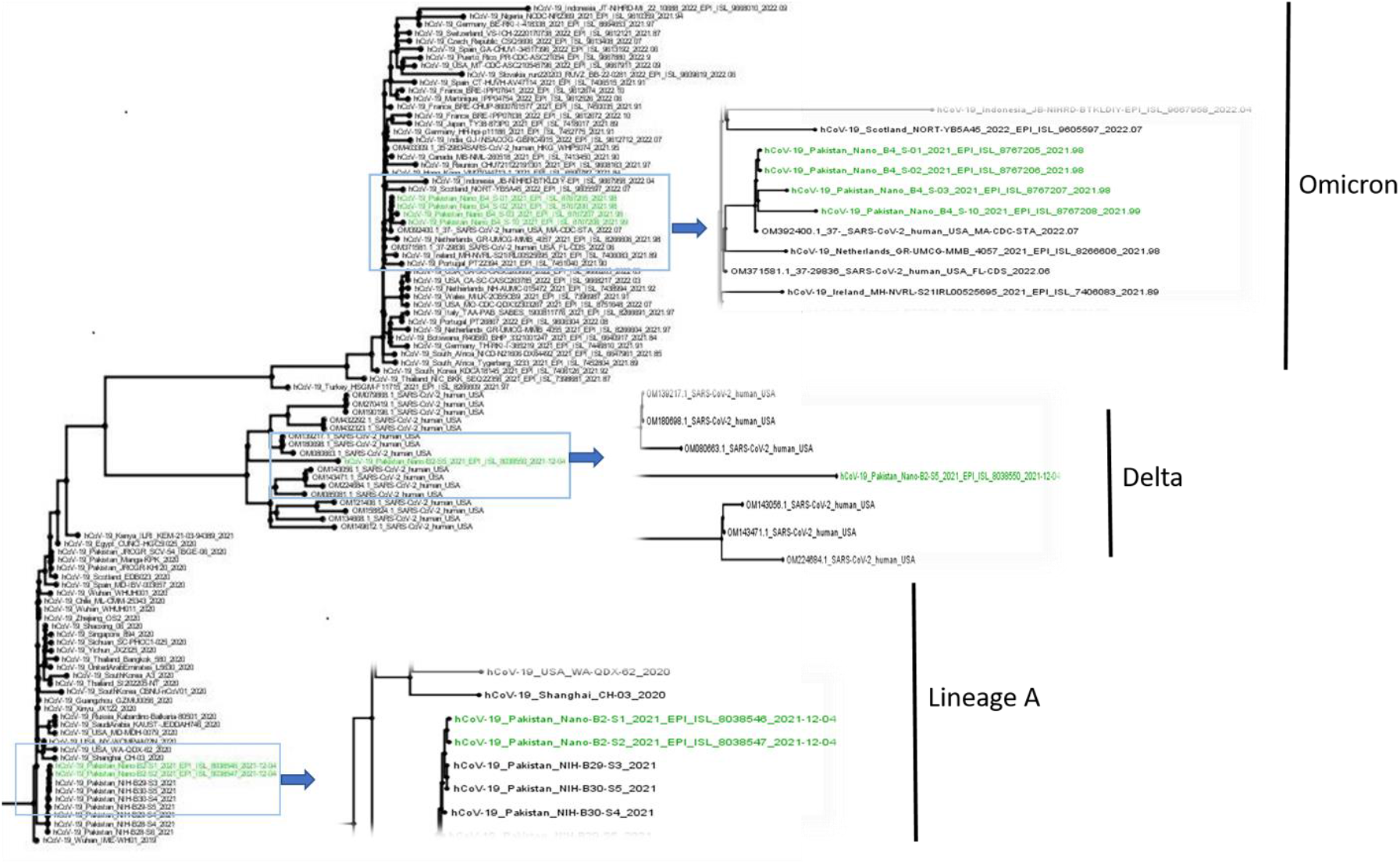
Phylogenetic tree of Omicron, Delta, and Lineage A variant viruses. Phylogenetic analysis was performed using the 108 full-length SARS-CoV-2 genomes from different pandemic countries obtained from the NCBI SARS-CoV-2 Resources (https://www.ncbi.nlm.nih.gov/sars-cov-2/) were subjected to Multiple Sequence Alignment (MSA) using MAFTT online server. The MSA was subsequently used to generate a Maximum Likelihood (ML) phylogenetic tree using IQtree (http://www.iqtree.org/). For reference, Wuhan SARS-CoV-2 (hCoV-19/Wuhan/IME-WH01/2019) sequence was used. The tree was edited and visualized using Figtree software (http://tree.bio.ed.ac.uk/software/figtree/).

### Mutation Profiling

The SARS-CoV-2 lineage diversity have been investigated in the current study. The study have shown the Delta (n=01) and Omicron (n=04) variant of SARS-CoV-2. Other than the omicron and delta variant, the study has also shown two lineage A isolates. Table 1 summarizes the mutation profile of all the study isolates. In case of omicron isolates, the BA.1 has been found to be the dominant lineage. While in case of delta variant, the sub lineage AY.27 has been identified. The mutational analysis of omicron variant has shown the characteristics mutations in the spike protein (A67V, H69-, V70-, T95I, G142-, V143-, Y144-, Y145D, N211-, L212I, G339D, S371L, S373P, S375F, K417N, N440K, G446S, S477N, T478K, E484A, Q493R, G496S, Q498R, N501Y, Y505H, T547K, D614G, H655Y, N679K, P681H, N764K, D796Y, N856K, Q954H, N969K). The spike is one of the most mutated protein in all the isolates. In ORF1a, four deletions (S2083-, L3674-, S3675-, and G3676-) and six substitutions (K856R, L2084I, A2710T, T3255I, P3395H, and I3758V). In case of nucleocapsid protein we have found a deletion of three amino acids (E31-, R32-, and S33-) and three substitutions (P13L, R203K, and G204R). In case of membrane glycoprotein only two substitutions (Q19E, A63T) were found. In case of ORF9b, three amino acid deletion (E27-, N28-, and A29-) and one substitution (P10S) was observed. No amino acid substitution and deletion has been observed in ORF3, ORF6-ORF8, ORF10 and Envelope glycoprotein.

**Table 1:** The mutation profile of seven study isolates

| GISAID ID | Lineage | ORF1ab | Spike | Membrane | Nucleocapsid | ORF3 | ORF8 | ORF9b |
| --- | --- | --- | --- | --- | --- | --- | --- | --- |
| EPI_ISL_8767205 | BA.1 | ORF1a: K856R, S2083-, L2084I, A2710T, T3255I, P3395H, L3674-, S3675-, G3676-, I3758V<br>ORF1b: P314L, I1566V | A67V, H69-, V70-, T95I, G142-, V143-, Y144-, Y145D, N211-, L212I, G339D, S371L, S373P, S375F, K417N, N440K, G446S, S477N, T478K, E484A, Q493R, G496S, Q498R, N501Y, Y505H, T547K, D614G, H655Y, N679K, P681H, N764K, D796Y, N856K, Q954H, N969K | Q19E, A63T | P13L, E31-, R32-, S33-, R203K, G204R |  |  | P10S, E27-, N28-, A29- |
| EPI_ISL_8767206 | BA.1 | ORF1a: K856R, S2083-, L2084I, A2710T, T3255I, P3395H, L3674-, S3675-, G3676-, I3758V<br>ORF1b: P314L, I1566V | A67V, H69-, V70-, T95I, G142-, V143-, Y144-, N211-, L212I, G339D, S371L, S373P, S375F, K417N, N440K, G446S, S477N, T478K, E484A, Q493R, G496S, Q498R, N501Y, Y505H, T547K, D614G, H655Y, N679K, P681H, N764K, D796Y, N856K, Q954H, N969K, L981F | Q19E A63T | P13L, E31-, R32-, S33-, R203K, G204R |  |  | P10S, E27-, N28-, A29- |
| EPI_ISL_8767207 | BA.1 | ORF1a: K856R, S2083-, L2084I, A2710T, T3255I, P3395H, L3674-, S3675-, G3676-, I3758V<br>ORF1b: P314L, I1566V | H69-, V70-, T95I, G142-, V143-, Y144-, Y145D, N211-, L212I, G339D, S371L, S373P, S375F, K417N, N440K, G446S, S477N, T478K, E484A, Q493R, G496S, Q498R, N501Y, Y505H, T547K, D614G, H655Y, N679K, P681H, N764K, D796Y, N856K, Q954H, N969K, L981F | Q19E A63T | P13L, E31-, R32-, S33-, R203K, G204R |  |  | P10S, E27-, N28-, A29- |
| EPI_ISL_8767208 | BA.1 | ORF1a: K856R, S2083-, L2084I, A2710T, T3255I, P3395H, L3674-, S3675-, G3676-, I3758V<br>ORF1b: P314L, I1566V | A67-, I68-, H69V, T95I, G142-, V143-, Y144-, N211-, L212I, G339D, S371L, S373P, S375F, K417N, N440K, G446S, S477N, T478K, E484A, Q493R, G496S, Q498R, N501Y, Y505H, T547K, D614G, H655Y, N679K, P681H, N764K, D796Y, N856K, Q954H, N969K | Q19E | P13L, E31-, R32-, S33-, R203K, G204R |  |  | P10S, E27-, N28-, A29- |
| EPI_ISL_8038546 | A |  | I68-, H69-, V70-, S71-, G72-, T73- |  |  |  | L84S |  |
|  |  |  | , N74-, G75-, T76- |  |  |  |  |  |
| EPI_ISL_8038547 | A |  | I68-, H69-, V70-, S71-, G72-, T73-, N74-, G75-, T76-, N679-, S680-, P681-, R682-, R683-, A684-, R685-, S686-, V687-, A688- |  |  |  | L84S |  |
| EPI_ISL_8038550 | AY.27 | ORF1a: A1306S, P2046L, P2287S, F2864L, V2930L, T3255I, T3646A, T4158I<br>ORF1b: P314L, Q348H, G662S, P1000L, M1596I, A1918V | G446V, L452R, T478K, D614G, D950N | I82T | R203M, G215C | S26L, G49V, T221K | D119-, F120- | R32L |

In case of delta variant, two mutations observed in spike protein as G446V, L452R and T478K, and D950N. In case of ORF1a, eight substitutions (A1306S, P2046L, P2287S, F2864L, V2930L, T3255I, T3646A, and T4158I) have been observed while in case of ORF1b, six substitutions (P314L, Q348H, G662S, P1000L, M1596I, and A1918V) have been observed. One mutation have been located in each membrane (I82T) and ORF9b (R32L). Two substitutions (R203M, G215C) have been observed in nucleocapsid and a two amino acid deletion in ORF8 (D119- and F120-). In ORF3a three mutations (S26L, G49V, and T221K) have been found.

Interestingly, the one lineage A isolate harbors a nine amino acid deletion in the spike protein spanning 68-76 amino acid region. The other lineage A isolate instead of having a nine amino acid deletion at amino terminal, another ten amino acid deletion has been observed at carboxy terminal spanning 679-688 amino acid region. These two isolates have only one substitution (L84S) in the ORF8 while all the other proteins are conserved.

## Discussion

During the first year of COVID-19 pandemic in the world, the SARS-Cov-2 constituted few mutations with the circulating viruses being closely related to Wuhan-HU-1 strain. The D614G substitution was the first mutation found in the SARS-CoV-2 with worldwide prevalence and now the D614G is one of the commonest mutation found in all the circulating lineages and sub-lineages of SARS-CoV-2 [4]. This mutation was found to be associated with increased transmissibility and binding affinity of virus with ACE2 compared to D614 viral strains. The dynamics of pandemic has changed globally after September, 2020 with the emergence of new viral strains carrying large number of mutations in the genome. These new variants has been characterized as alpha, beta, gamma, and delta. Despite these major lineages of the SARS-CoV-2, there are many other lineages and sub-lineages of virus has emerged that changed the global pattern of pandemic. These variants gives the virus a survival advantage by increasing transmissibility, infectivity and immune escape from neutralizing antibodies [10, 11].

Although restrictive measures on limiting international travel and local lockdown on key hotspot areas have been in practice from time to time, the variants still find their way of emergence [12]. Similar scenario was observed in Pakistan, where the first case of SARS-CoV-2 was detected from a traveler. Like all parts of the World, the Pakistan has also been hit by the pandemic in 2020 with Wuhan-HU-1-like strains of SARS-CoV-2. The first and second pandemic wave in Pakistan has been led by lineages of SARS-CoV-2 of the Wuhan-HU-1 strain. While the third wave of pandemic was led by different variants of SARS-CoV-2. Among these, alpha, beta and delta was found to be the dominant one. After the third wave, the alpha and beta strains were not apparent and fourth wave in Pakistan was taken over by predominantly delta variant [13]. Pakistan witnessed almost four months long fourth wave of pandemic starting from July, 2021 and ended in mid of October, 2021. The genomic surveillance has revealed the prevalence of delta variant during fourth pandemic wave but the data of circulating lineages after the fourth wave was unavailable. The current study has identified the lineages other than delta in the population of Islamabad during the flattened curve achieved after the fourth wave. We have found three different lineages during the month of November and December, 2021. The drop in the number of cases after October, 2021 may be attributed to the circulation of different lineages in the population. The other contributing factor is the increase in vaccination status of individuals. As of January 24, 2022, 35.7% of population has been fully vaccinated. Another important contributing factor may be the circulation of lineage A in the population. The original Wuhan-Hu-1 strain belonged to the lineage A and later the mutations in the lineage A led to the emergence of lineage B and other lineages. The circulation of lineage A in the population may also one of the reason of this flattened curve. The first two sequences of lineage A were reported in June, 2020 and then in April, and May, 2021 two more sequences of lineage A were reported from Pakistan. Interestingly, during November and December, 2021 a total of nine whole genomes of lineage A were reported. One of the study isolate of Lineage A harbors a ten amino acid deletion in the furin cleavage site. Studies have reported that deletion of this site may impair the viral pathogenicity but this need to be validated further [14].

The omicron variant harbors more than 30 mutations in the spike glycoprotein with almost 15 mutations in the receptor binding domain (319-541 amino acids). These mutations may weaken the vaccine protection in the individuals. This is one of the rapidly spreading variant of SARS-CoV-2 and now it is becoming the dominant lineage in the countries where it is first time detected. As observed in South Africa, UK, USA, Germany, Scotland, Belgium, Denmark, and France. Omicron multiplies 70 times more in the lung airways as compared to delta but not affect deep into the lungs [14]. Hence, it is less severe than delta.

In Pakistan the first case was reported in 09 December, 2021 and since then the number of cases have been increasing gradually. During late December, 2021 the government of Pakistan announced the commencement of fifth COVID-19 wave in Pakistan. Since that the number of cases start increasing and currently there have been 7,195 positive cases in a single day with positivity rate of 12.5% (as of January 24, 2022). The increase in number of cases demands for enhancement of the vaccination of population, imposition of the protective measures like maintain social distance, wear mask, avoid social gathering.

To conclude, the SARS CoV2 spread pattern has been no different with worldwide comparison, with a first and second wave of pandemic being dominated by WUHAN-HU-1 like strains while the third and fourth wave was led by different variants of concern that lead to increase in number of infections and deaths. The emergence of new variants of SARS-CoV-2 demands for increasing the genomic surveillance in country in order to track the emergence and spread of new variants of SARS-CoV-2.

## Data Availability

The sequences generated in the study were submitted to GISAID (https://www.gisaid.org/) with accession IDs: EPI_ISL_8038546, EPI_ISL_8038547, EPI_ISL_8038550, EPI_ISL_8767205, EPI_ISL_8767206, EPI_ISL_8767207, EPI_ISL_8767208.

